# A click chemistry-based biorthogonal approach for the detection and identification of protein lysine malonylation for osteoarthritis research

**DOI:** 10.1101/2024.12.12.628274

**Authors:** Anupama Binoy, Pandurangan Nanjan, Kavya Chellamuthu, Huanhuan Liu, Shouan Zhu

## Abstract

Lysine malonylation is a post-translational modification where a malonyl group, characterized by a negatively charged carboxylate, is covalently attached to the ε-amino side chain of lysine, influencing protein structure and function. Our laboratory identified Mak upregulation in cartilage under aging and obesity, contributing to osteoarthritis (OA). Current antibody-based detection methods face limitations in identifying Mak targets. Here, we introduce an alkyne-functionalized probe, MA-diyne, which metabolically incorporates into proteins, enabling copper(I) ion-catalyzed click reactions to conjugate labeled proteins with azide-based fluorescent dyes or affinity purification tags. In-gel fluorescence confirms MA-diyne incorporation into proteins across various cell types and species, including mouse chondrocytes, adipocytes, Hek293T cells, and *C. elegans*. Pull-down experiments identified known Mak proteins such as GAPDH and Aldolase. The extent of MA-diyne modification was higher in Sirtuin 5-deficient cells suggesting these modified proteins are Sirtuin 5 substrates. Pulse-chase experiments confirmed the dynamic nature of protein malonylation. Quantitative proteomics identified 1136 proteins corresponding to 8903 peptides with 429 proteins showing 1-fold increase in labeled group. Sirtuin 5 regulated 374 of these proteins. Pull down of newly identified proteins such as β-actin and Stat3 was also done. This study highlights MA-diyne as a powerful chemical tool to investigate the molecular targets and functions of lysine malonylation in OA conditions.

## 1. Introduction

Post-translational modification of proteins refers to the biochemical covalent, enzymatic, and non-enzymatic addition of functional groups on proteins following their synthesis from ribosomes^1^. These functional groups can be small electrophilic biochemical metabolites like phosphate, sugars, nucleotides, methyl group, and acetyl group, long chain/short chain acyl chains like palmitoyl, malonyl groups, radicals generated from redox reactions like *S*-nitrosylation (SNO) and *S*-glutathionylation etc^2^. Post-translational modifications induce changes in amino acid chemical properties such as deamination, deamidation, citrullination, and oxidation, as well as protein autocatalysis that results in protein backbone cleavage. It offers complex and diverse functional roles to the existing proteome by regulating protein activity, structure and conformation, its location, and molecular interactions^3^. Post-translational covalent modification of proteins is fundamental to numerous cellular and biological functions. It plays a significant role in regulating normal cell physiology as well as the pathogenesis of diseases^4^. Lysine malonylation (Mak) is a reversible protein post-translational modification wherein a malonyl group is added to the ε-amino group of the lysine residue in a protein. The addition of a negatively charged carboxylate group to the protein imparts significant changes in the protein’s structure and function^5^. The reversible protein malonylation is theoretically regulated by acyltransferases and deacylases. The enzymes responsible for catalyzing the transfer of a malonyl group remain largely unidentified. However, recent work by Zang et al. has provided evidence that KAT2A is involved in histone malonylation^6^. Meanwhile, Sirtuin 5 (Sirt5), a class III lysine deacetylases member, is known for its lysine demalonylation activity along with its lysine desuccinylation function^7,8^. Our lab has previously demonstrated that deficiency of Sirt5 in mouse primary chondrocytes increases the global protein MaK level and disrupts cellular metabolism^8^. Identification of the protein substrates of Sirt5 is crucial for further elucidating the function of malonylation on the cellular metabolism and pinpoint the downstream targets. The anti-malonyl lysine antibody has been used previously to enrich and identify several malonylated proteins^8-10^. Antibody-based enrichment is generally based on the development of different antibody domains specific to targeting antigens with specific PTM, followed by immunoaffinity pull-down of target peptides from the protein lysates. However, this strategy suffers from a lack of specificity by missing some of the target PTM site due to the presence of adjacent PTM present on the same sequence^11^. To overcome this challenge, we took a chemical approach, utilizing an alkyne functionalized chemical probe to efficiently detect and quantify protein substrates of lysine malonylation through fluorescence visualization and further identify them using a quantitative proteomics approach (Figure 1B). Alkyne functionalized probe allows metabolic detection of protein substrates by utilizing the copper(I) ion-catalyzed alkyne azide cycloaddition click chemistry to conjugate the labeled proteins with azide-fluorescent dyes or affinity purification tags^12^. This approach has been widely utilized in identifying several other types of PTMs including lipidation like myristiolation^13,14^, succinylation^15,16^, as well as acetylation^17,18^ and glycosylation^19,20^. Previously, Bao et al. developed a malonic acid-based chemical probe called MalAM-yne to detect lysine malonylation^21^. In this study, we designed and synthesized a novel Meldrum’s acid-based probe, i.e. MA-diyne, as a better synthetic alternative to the previously reported probe^21^. Meldrum’s acid or isopropylidene malonate is a condensation product of malonic acid and acetone. We developed this new probe by functionalizing Meldrum acid or isopropylidene malonate with alkynylation under control conditions (Figure 1A). We hypothesize that Meldrum acid will be readily absorbed into the cells due to its cyclic structure and subsequently will get linearized to form a malonyl group by an unknown action of intracellular esterases, thus leading to effective detection and identification of the malonylated proteins.

**Figure 1:**
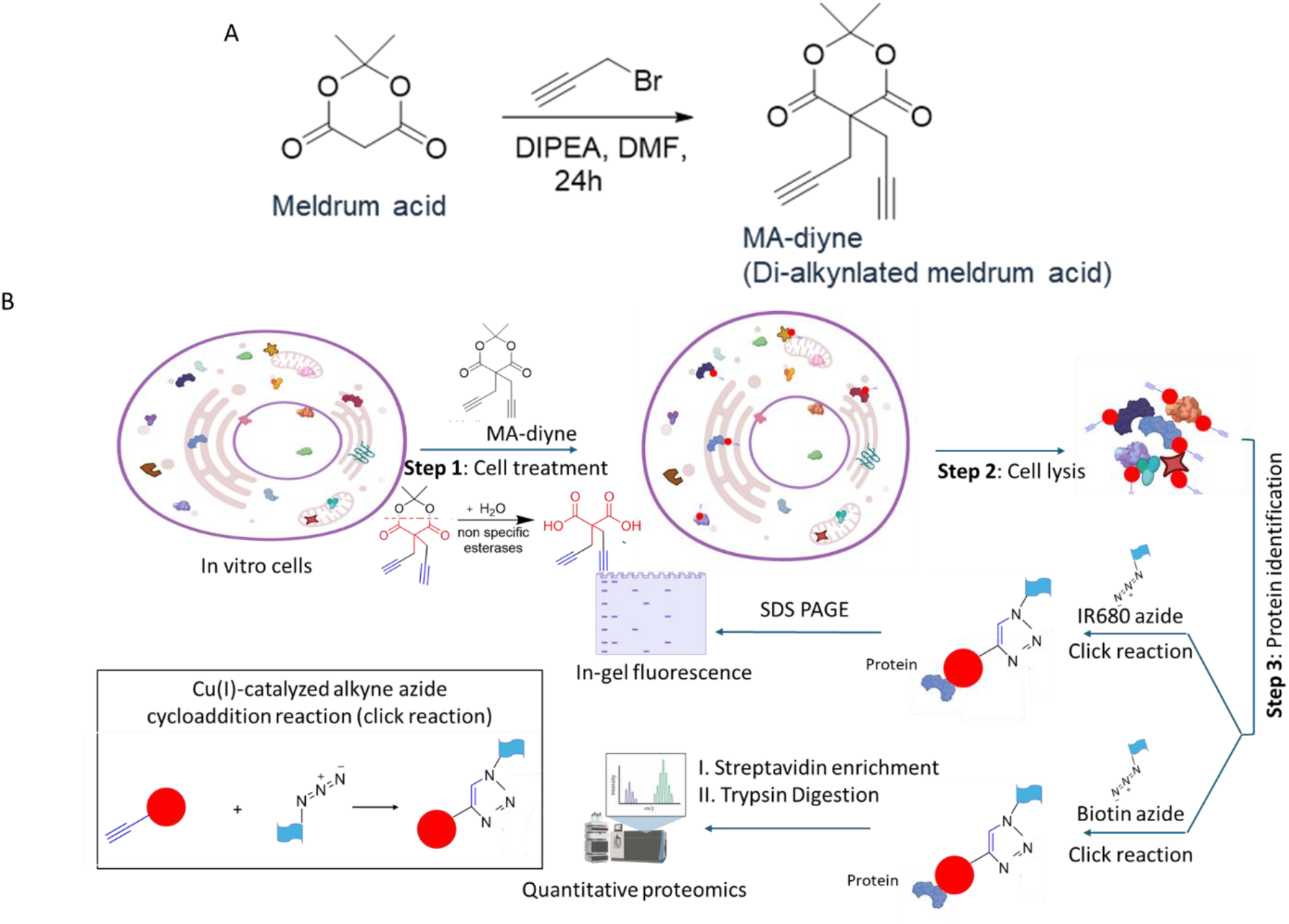
A. Design, synthesis, and characterization of MA-diyne. B. Schematic description of the workflow to detect and identify lysine malonylated proteins using MA-diyne.

## 2. Results and discussion

### 2.1. Chemical synthesis

Bao et al 2013 were the first to report a chemical probe with an alkyne handle on the malonate group named Mal-yne^21^. However, they modified this probe to increase cell permeability by masking the carboxylate sides with two acetoxymethyl groups and called it MalAM-yne. In this report, we selected an alternative approach by adding an alkyne handle to Meldrum acid or isopropylidene malonate. The high reactivity of this molecule is attributed to the methylene group present between the carbonyl group and this site was utilized to add the propargyl unit to meldrum acid under controlled conditions. Thereby, we generated dipropargyl meldrum acid or MA-diyne through the following chemical synthesis (Figure 1A). The proposed probe has a better ability of hydrolysis in the presence of esterase than the probe MalAM-yne by Bao et al 2013^21^. The synthesis involves di-alkynylation on commercially available Meldrum acid. The reaction is performed under mild conditions such as DIPEA (diisopropyl ethylamine) with anhydrous DMF solvent under room temperature. The reaction is stirred for 24 hr. After the working up of the reaction, the product is taken for purification. The initial purification of the MA-diyne product is carried out with normal SiO_2_ which leads to ring-opening of MA-diyne due to its acidity. Later, SiO_2_ was neutralized with triethylamine and then purification was carried out. The pure product is characterized by ^1^H NMR spectroscopy [^1^H NMR (400 MHz): δ 1.84 (s, 6H), 2.18, (t, *J* = 5.2 Hz, 2H), 2.88 (d, *J* = 2.8 Hz, 4H) ppm. The respective assignments and full spectra of MA-diyne are given as supplementary information (Scheme SF1 and SF2). The described method (one step) is simple, faster, and gives maximum yield (40%) compared to the previously reported three-step method (overall yield; 23%). The yield limitation to 40% in the present method arises from the ring-opening reaction of Meldrum’s acid under the specified conditions.

### 2.2 MA-diyne can be metabolically incorporated into cellular proteins

Our first inquest was to analyze whether MA-diyne can metabolically incorporate into proteins similarly to malonyl-CoA. Therefore, we incubated mouse primary chondrocytes with different concentrations (0-200 µM) of MA-diyne in complete medium for 6 h (a working stock of 100 mM was prepared by dissolving MA-diyne in DMSO; DMSO was taken as control). The protein lysate was collected after harvesting the cells and was subjected to azide-alkyne click chemistry to conjugate the alkyne side group with IR680 azide dye. The clicked proteins were then resolved on SDS-PAGE and scanned using in-gel IR fluorescence imager. The result showed that there was a dose-dependent increase of MA-diyne labeling of global proteins with the optimal concentration at less than 50 µM MA-diyne (Figure 2A). To analyze the time required for the metabolic labeling, a time-dependent experiment was conducted by incubating the primary chondrocytes with 100 µM of MA-diyne at different time points ranging from 0-12 h. MA-diyne was able to efficiently label the proteins in no more than 2 h (Figure 2B). Interestingly, we observed that MA-diyne showed more bands in gel fluorescence compared to western blot analysis of the same proteins from primary chondrocytes using a commercial anti-KMal antibody (Supplementary Figure 1). The dose and time-dependent experiment demonstrated the efficient labeling of proteins by MA-diyne.

**Figure 2:**
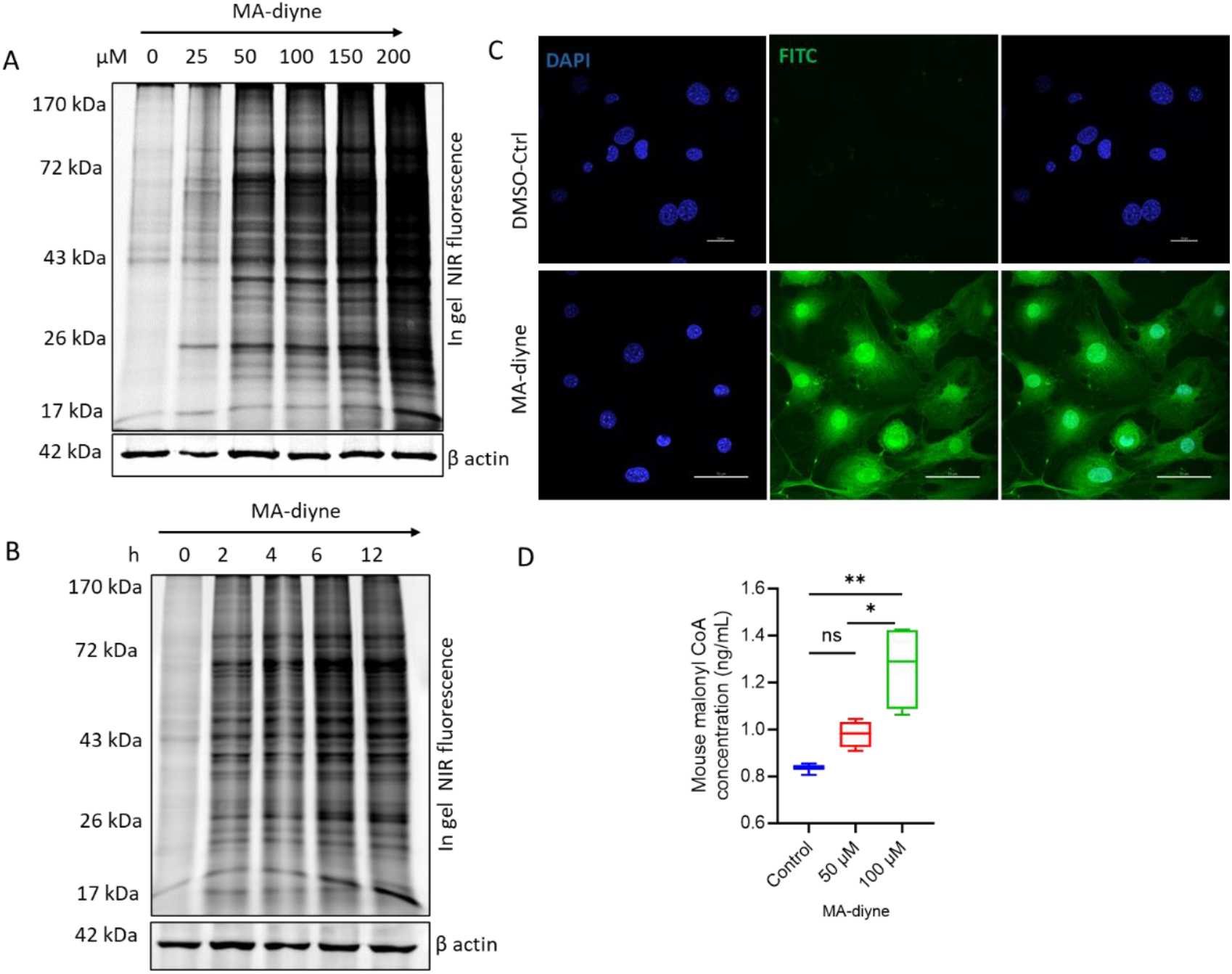
Assessment of the ability of MA-diyne to metabolically label proteins. A. Mouse primary chondrocytes were incubated with the indicated concentration of MA-diyne for 6 h. The cell lysates were then clicked with IR680-azide followed by in-gel fluorescence analysis. B. Mouse primary chondrocytes were incubated with 100 µM of MA-diyne for the indicated time points. Cell lysates were then clicked with IR680-azide and in-gel fluorescence analyses. β actin was used as a loading control. C. Confocal microscopic fluorescence image depicting the ready uptake of the MA-diyne into the cells and subcellular localization of malonylated proteins in the primary chondrocytes. D. Box plot showing the concentration of intracellular malonyl-CoA in the cells after incubation with different concentrations of MA-diyne for 4 h. n=4. Data are presented as mean ± SEM. Three group comparisons were evaluated using two-way ANOVA. Significance is noted as ns p>0.05, *p<0.05, and **p<0.01.

To confirm that the signal achieved in the primary chondrocytes after treatment with MA-diyne is due to metabolic labeling and not due to nonspecific binding, we used qualitative fluorescence imaging to visualize the labeled proteins. The primary chondrocytes were cultured on coverslips overnight and treated with different doses of MA-diyne for 6 h. The cells were immediately fixed using ice-cold 4% paraformaldehyde and then permeabilized. The cells were then subjected to click reaction using Carboxyrhodamine 110 Azide for 1 h. The cells on the coverslips were mounted on clean glass slides using mounting reagent with DAPI. Control samples were generated by performing the same procedure without MA-diyne or Carboxyrhodamine 110 Azide. The slides were imaged using Nikon A1R confocal microscopy under 60X magnification (Figure 2C and supplementary Figure 2). MA-diyne was found to be rapidly absorbed into the cells which was demonstrated by the intensity of labeling only in the samples treated with MA-diyne followed by click reaction with Carboxyrhodamine 110 Azide but not in the control samples incubated with Carboxyrhodamine 110 Azide only or samples treated with only MA-diyne (supplementary Figure 2). We observed widespread protein labeling in various cell compartments, prompting us to conduct quantitative proteomics. However, to ensure the labeling is due to the malonyl-CoA formed by the acyclization of Meldrum acid, we estimated the concentration of malonyl-CoA formed in the cells after incubating the cells with MA-diyne in different concentrations and compared it with control cells without MA-diyne. We observed a significant increase in the concentration of malonyl CoA at 100 µM. However, as protein labeling by MA-diyne increases with incubation time, malonyl-CoA concentration may also depend on the time needed for MA-diyne uptake and linearization.

### 2.3 MA-diyne was dynamically removed from the proteins representing the reversible nature of lysine malonylation

Lysine modifications like succinylation, glutarylation, and malonylation are reversible forms of protein post-translational modifications^9,22,23^. Although the enzyme responsible for adding a malonyl group to the proteins is still unknown, Sirtuin 5 (Sirt5) has been well-documented as the enzyme responsible for removing this modification^6,24^. To examine whether the labeling by MA-diyne is also reversible, we conducted a pulse-chase experiment. The wild type and Sirt5 knockdown primary chondrocytes were first incubated with 200 µM MA-diyne for 1 h. The cells were then washed and chased with 200 µM Meldrum acid for 0-5 h. The cells were harvested at different time points, and protein lysates were conjugated with IR680-azide dye. The in-gel fluorescence imaging reveals that the labeling signal by MA-diyne in wild-type primary chondrocytes started fading after 0.5 h of Meldrum acid incubation, indicating the removal of MA-diyne labeling from the proteins (Figure 3A). In comparison, in Sirt5 knockdown cells, the labeling signal with MA-diyne became more intense after 0.5 h compared to the baseline level and there is less removal of MA-diyne from the labeled proteins compared to the wild-type cells (Figure 3B). We also noticed that the labeling by MA-diyne was not modulated by the endogenous levels of malonyl-CoA, evidenced by no difference between wild type and cells deficient of acetyl-CoA carboxylase (ACC1), an enzyme that produces malonyl-CoA (supplementary figure 3). These results suggested that the metabolically modified proteins by MA-diyne could be substrates of Sirt5 demalonylase and the process is reversible.

**Figure 3:**
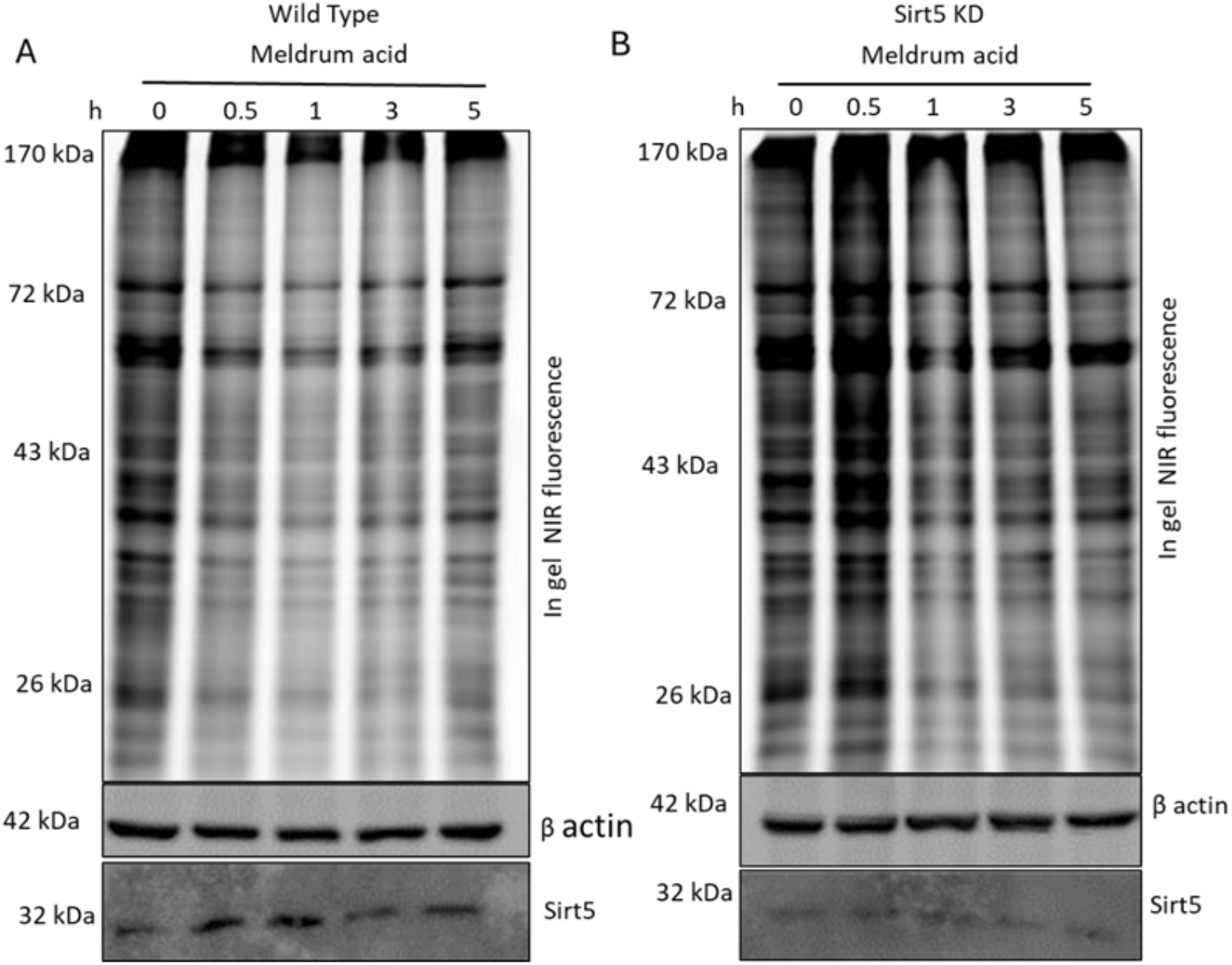
Pulse-chase experiment to determine the dynamic nature of lysine malonylation. Wild-type mouse primary chondrocytes (A) and Sirt5KO primary chondrocytes (B) were labeled with 200 µM MA-diyne for 1 h and then pulse chased with 200 µM Meldrum acid (precursor). The lysates were collected at the indicated time points followed by a click reaction with IR680-azide and in gel fluorescence analysis. β actin was used as a loading control. The same blot was probed with an anti-Sirt5 antibody to quantify the knockdown efficiency.

### 2.4 MA-diyne successfully identified putative malonylated proteins

Our previous studies have demonstrated the important role of Sirt5 in chondrocyte metabolism by regulating the malonylation of metabolic enzymes^25-28^. We have also used proteomics to identify the malonylome in chondrocytes using the traditional Kmal PTMscan antibody-based enrichment method^28^. We reported the enrichment of 1000 peptides corresponding to 469 proteins. Herein this report, we applied a chemoproteomics-based approach to enrich malonylated proteins from primary chondrocytes using MA-diyne. Wild type and Sirt5 knockdown primary chondrocytes were labeled with MA-diyne (100 µM) for 6 h and then the protein lysates were conjugated to biotin azide through azide-alkyne copper cycloaddition reaction. The biotin-conjugated proteins were then immuno-precipitated using avidin agarose beads. The beads were thoroughly washed with HPLC water to prevent detergent or protease inhibitor contamination in the LCMS, followed by addition of 9M urea wash. The beads were then subjected to reductive alkylation with dithiothreitol (DTT) and iodoacetamide (IAA), on bead trypsin digestion, and desalting. The enriched peptides were then subjected to bottom-up quantitative proteomics on Orbitrap Astral Instrument. A total of 1136 proteins corresponding to 8903 peptides across all samples (supplementary data excel 1) were identified, which was 2.4 times more than the proteins we identified before using Kmal PTMscan antibody enrichment^28^. 430 proteins were seen to show a more than 1-fold increase in probe group in comparison to the control group with more than 6 unique peptides (supplementary data excel 2). This indicated that MA-diyne could enrich malonylated proteins. The enrichment of proteins in the control group accounts for the endogenously biotinylated proteins which might have been enriched with avidin beads. Furthermore, 387 out of the 430 proteins were found to have more than 1-fold increase in the Sirt5 knockdown + MA-diyne group in comparison to the wild type + MA-diyne group (supplementary data excel 2). This indicates that Ma-diyne successfully enriched proteins which are regulated by Sirt5 demalonylase enzyme. Since the peptides search was not done for the modified peptides (due to the difficulty in eluting the biotinylated peptides after on-bead trypsinization), we compared the 8903 peptides to those modified sites identified by Kmal PTMscan antibody enrichment in our previous study. Interestingly, some of the peptides sequences identified in MA-diyne enrichment matched with the sequence identified in the LC-MS/MS data acquired after enrichment with the Kmal PTMscan antibody^28^ (Table 1). All these observations indicate that MA-diyne can efficiently detect and identify lysine malonylated proteins. Since our data lack site-specific identification of malonylation, our lab is currently exploring alternative proteomic approaches to identify malonylated sites on proteins detected by MA-diyne in MA-diyne treated primary chondrocytes. Further, these proteins will be pulled down using enrichment tags and analyzed in LC-MS/MS for the signature satellite peaks.

**Table 1:**
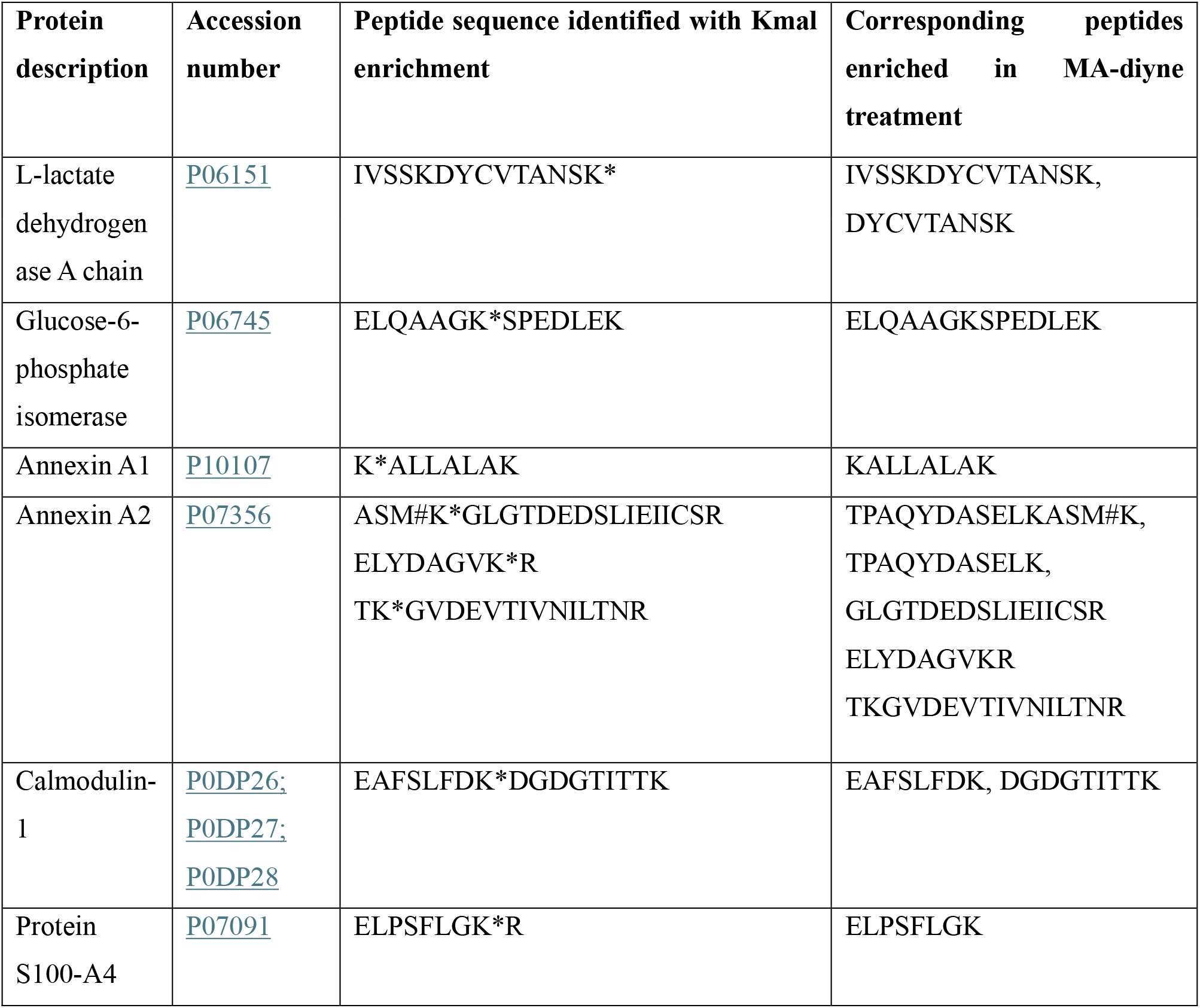
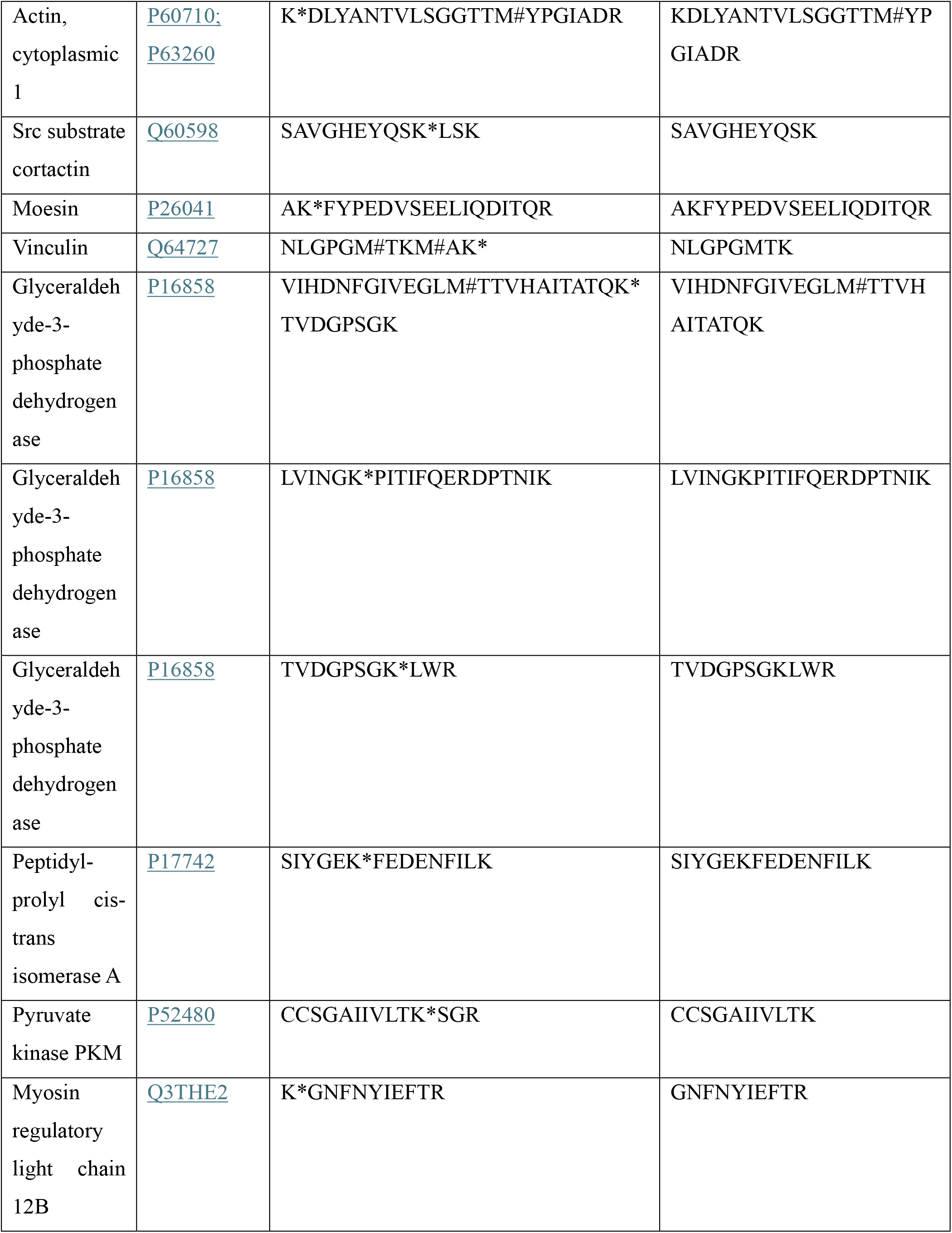
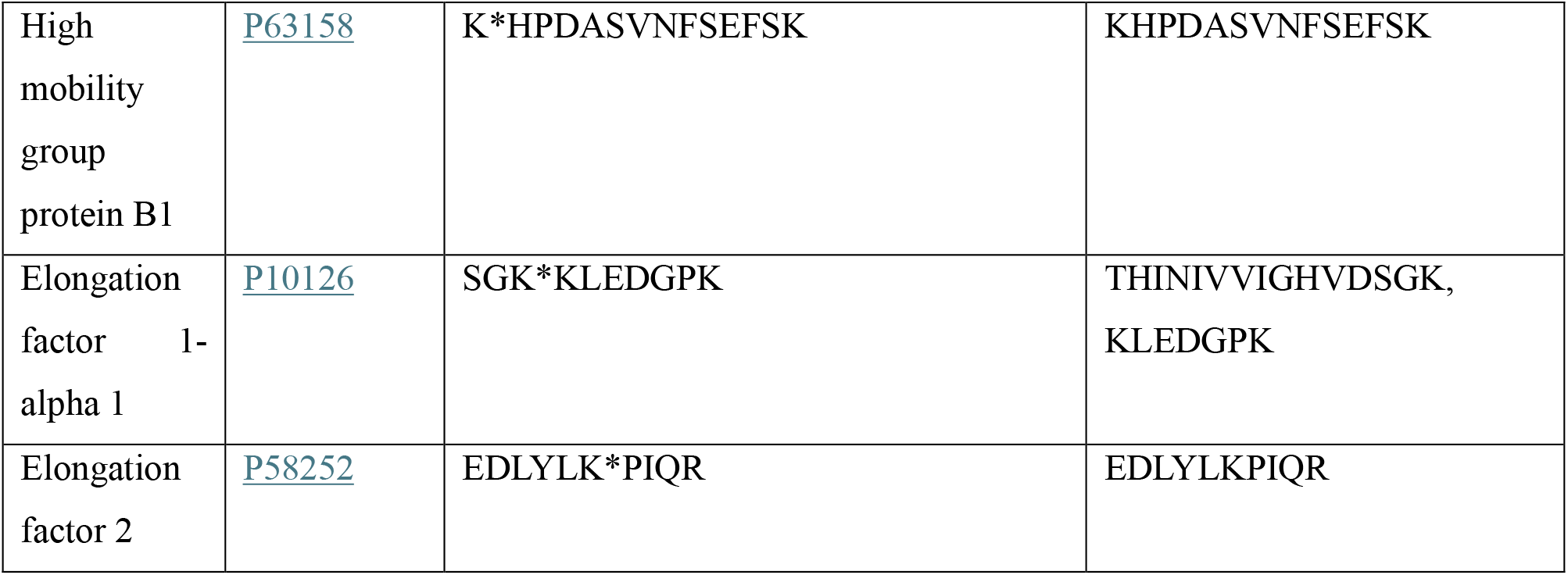
List of manually validated peptides sequences with modified malonylated sites.

We then conducted a pull-down experiment to validate further that MA-diyne could enrich malonylated proteins. The eluted protein from the avidin beads in a separate experiment were resolved on the SDS-PAGE followed by immunoblotting against antibodies for some already known malonylated proteins, ALDOA (Fructose-bisphosphate aldolase A)^28^ and GAPDH (Glyceraldehyde-3-phosphate dehydrogenase)^29^. The detection of GAPDH and ALDOA by western blot analysis of the eluted proteins from the beads (Figure 4A) confirmed that MA-diyne can indeed enrich malonylated proteins. We could also pull down other novel proteins in our proteomics list like Stat3, and β actin that haven’t been reported before (Figure 4B).

**Fig 4.**
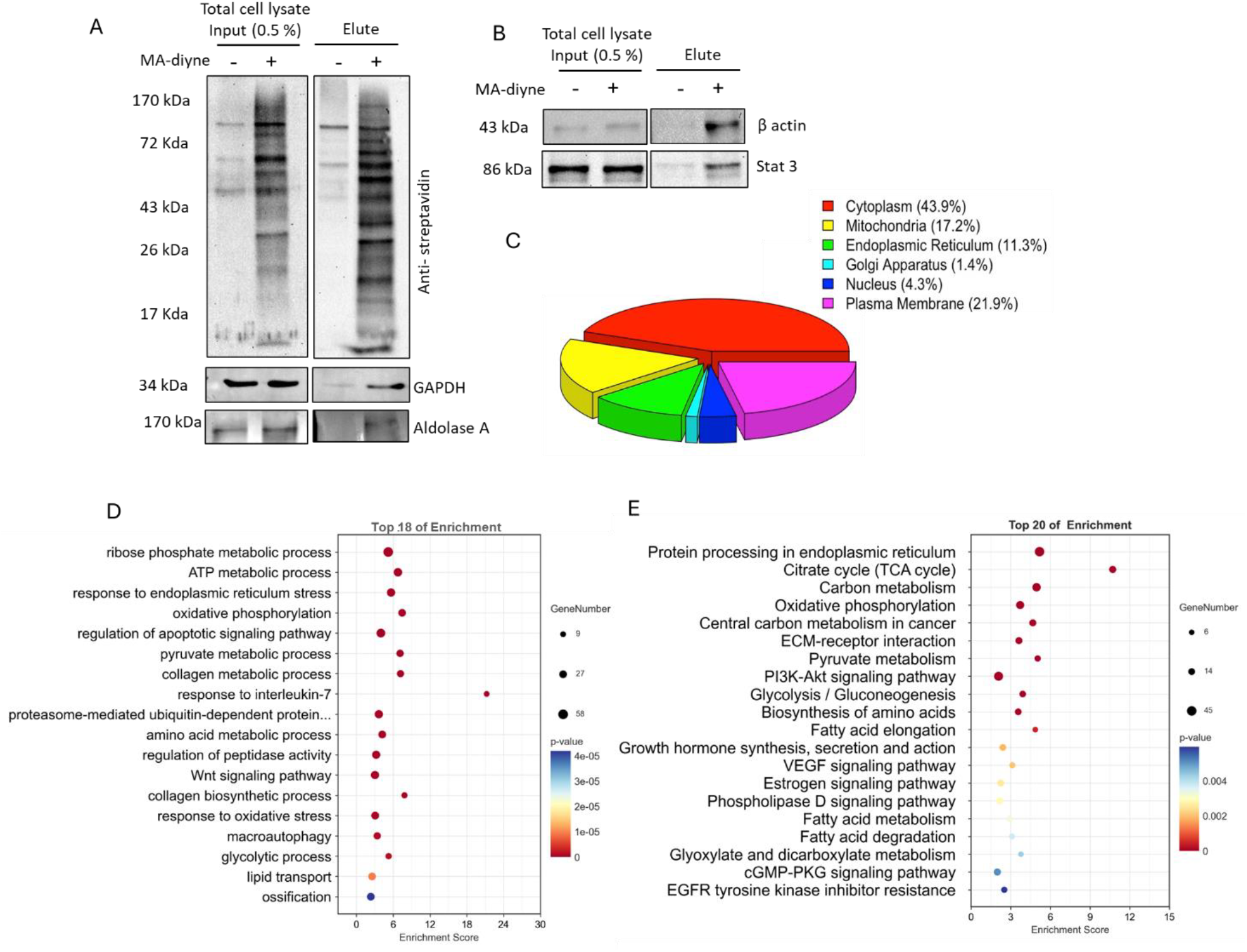
Validation of Lysine malonylated protein targets of MA-diyne. A. Pull down of known malonylated proteins such as GAPDH and Aldolase A after enrichment using avidin beads. n=2. B. pull down of newly identified proteins after MA-diyne enrichment. n=2. C. Pie chart depicting the percentage of malonylated proteins found in different subcellular compartments by GO analysis. D. distribution of enriched proteins based on their biological process by GO analysis, E. KEGG pathway analysis of the enriched proteins. p value < 0.05.

Gene ontology (GO) and pathway analysis of the identified proteins revealed that the malonylated proteins are localized differentially in various subcellular compartments including cytoplasm, plasma membranes, endoplasmic reticulum, nucleus, mitochondria, and Golgi apparatus of the primary chondrocytes (Figure 4C and supplementary excel 3). These findings align with similar subcellular localization patterns of malonylated proteins observed in other cell types, such as plants^30^, prokaryotes^31^, parasites^32^ and mammalian cells^9^. GO analysis based on the biological processes revealed that the identified malonylated proteins belonged to processes related to musculoskeletal tissues like collagen biosynthesis, and ossification. Additionally, it was revealed that malonylated proteins are highly involved in glucose, amino acid, and pyruvate metabolic and lipid transport processes. The identified proteins are also enriched in biological processes like ribose phosphate metabolism, ATP production, TCA cycle, glycolysis, and oxidative phosphorylation (Figure 4D and supplementary excel 4). These findings are consistent with previous reports on the involvement of lysine malonylation in regulation of metabolic disorders like type 2 diabetes^33^, cardiovascular diseases^34^, osteoarthritis^25,35^ etc. We also observed the role of malonylated proteins in the mechanisms related to the quality control of proteins such as response to endoplasmic reticulum stress^5^, autophagy^36^ and proteasome-mediated protein processing (Figure 4D and supplementary excel 4). KEGG analysis of the identified malonylated proteins revealed regulation of several signaling pathways like VEGF, Wnt signaling, cGMP-PKG, PI3K-AKT, estrogen signaling, EGFR tyrosine kinase inhibitor resistance, growth hormone synthesis and secretion, phospholipase D signaling pathways etc (Figure 4E and supplementary excel 5).

### 2.5 MA-diyne can detect lysine malonylation in a wide range of cell types *in vitro*

To assess the applicability of MA-diyne probe in detecting malonylated proteins in different types of cells, we analyzed the metabolic labeling profile of MA-diyne in subcutaneous primary adipocytes (Figure 5A), and Hek 293T cell lines (Figure 5B). Both cells exhibited robust MA-diyne signaling compared to the control suggesting successful metabolic incorporation of MA-diyne into cellular proteins. In addition, we assessed whether MA-diyne can be used to metabolically label proteins of *C*.*elegans*. We incubated live *C*.*elegans* with 1mM MA-diyne for 6 h with constant shaking and then lysed the worms to extract proteins. The proteins were then conjugated with IR680 azide. Ingel fluorescence analysis revealed that MA-diyne resulted in a robust labeling of malonylated proteins (Figure 5C). The route for the metabolic labeling is unclear but it is assumed that MA-diyne could have penetrated *C*.*elegans* cells either via the esophageal route or epidermal absorption/passive diffusion.

**Figure 5:**
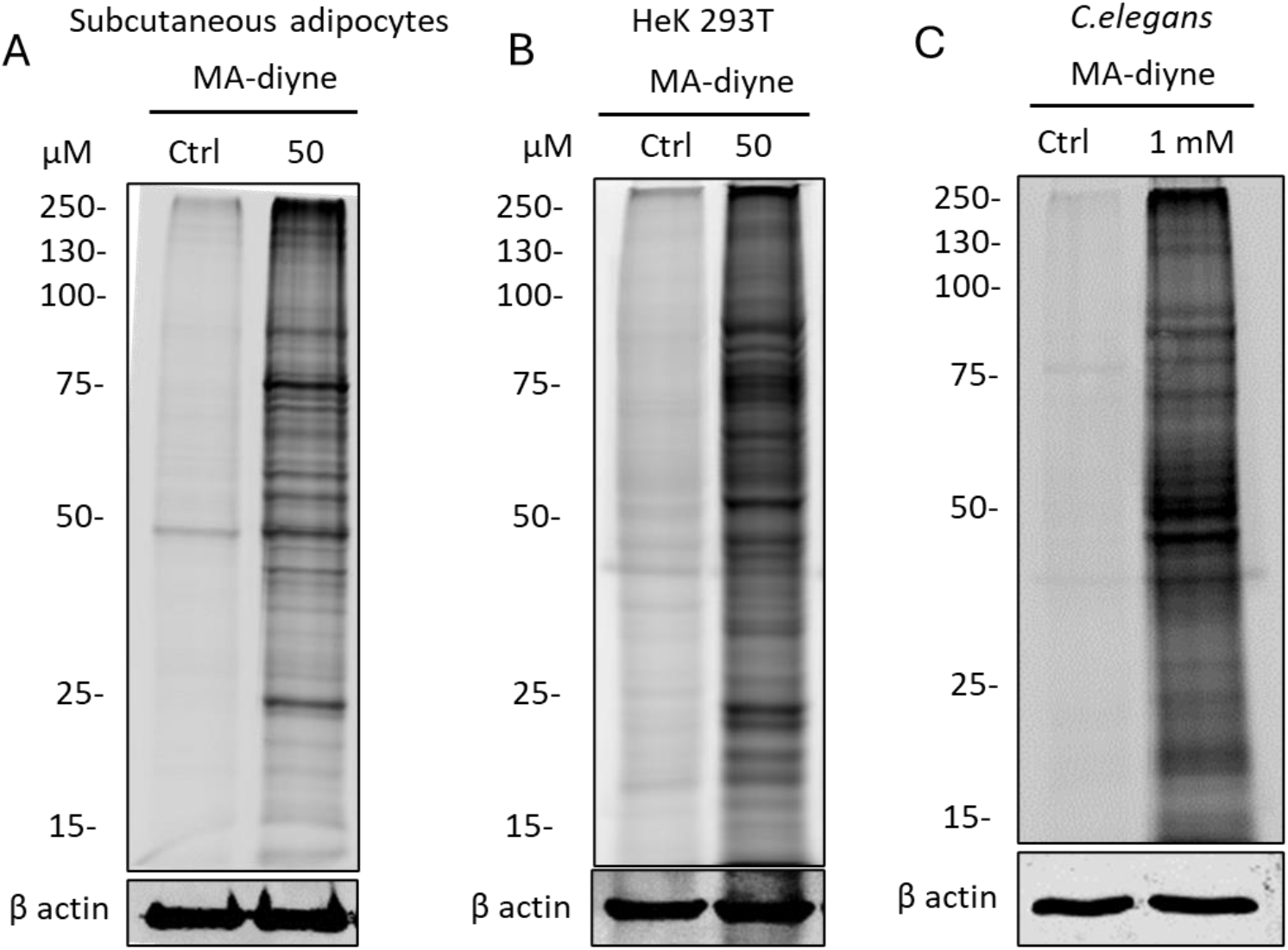
MA-diyne can detect lysine malonylated proteins in other cells like primary subcutaneous adipocytes, Hek293T cells and *C. elegans*, n=2

## 3. Conclusion

In summary, we have developed a novel chemical probe, MA-diyne for identifying and quantifying the protein malonylation in primary chondrocytes and several other types of cells. This probe was synthesized by adding a propargyl group to the meldrum acid for utilizing the conventional chemoproteomic approach to pull down the malonylated proteins. MA-diyne was observed to be readily uptaken into the cells and after getting acylized to malonyl-CoA by the action of intracellular esterases, it then metabolically labeled the proteins. It was also observed that labeling by MA-diyne was dynamic and regulated by the demalonylase enzyme, Sirt5. Moreover, the labeling by MA-diyne was observed to be nonenzymatic and not dependent on the endogenous levels of malonyl CoA. Quantitative proteomics could identify a significantly larger number of proteins than our previous attempt with the Kmal PTMscan antibody. Moreover, this further enables us to validate the enzymatic function of several identified metabolic proteins and their role in the progression of osteoarthritis.

## Acknowledgements

We thank the following funding support: Hevolution Foundation AGE award AGE-008 (SZ, HL), National Institutes of Health grant R01AR081804 (SZ, HL), National Institutes of Health grant R15AR080813 (SZ, HL), Arthritis National Research Foundation, American Society for Bone and Mineral Research FIRST award (SZ), Rheumatology Research Foundation Innovative Award (SZ), Osteopathic Heritage Foundation Ralph S. Licklider, D.O. Endowed Professorship (SZ). We thank the staff (including Tammy Mace, Angela Smith, Shawn Rosensteel, and Scott Carpenter) at the animal facility in the Life Science Building for their excellent care for our animals. We thank Dr. Vishwajeet Puri, Dr. Craig Nunemaker, and Dr. Kevin Lee for providing the cell lines. We also thank Dr. Nathaniel Szewczyk for helping us conduct the labeling in *C*.*elegans*. We acknowledge the help from Proteomics Core Facility at the Cell Signaling Technology Company for their help with the proteomics assay.

