## Supplementary Materials for "A click chemistry-based biorthogonal approach for the detection and identification of protein lysine malonylation for osteoarthritis research"

### Supporting Information

#### 1. General methods and materials

**Preparation of MA-diyne**: To a stirred solution of meldrum acid (1.0 g, 6.94 mmol, 1.0 eq.) in DMF (10 mL) was added DIPEA (6.04 mL, 34.7 mmol, 5.0 eq.) followed by Propargyl bromide (0.79 mL, 10.41 mmol, 1.5 eq.) at room temperature under inert atmosphere. The resulting reaction mixture was allowed to stir at room temperature for 24 h. The reaction mixture was then diluted with cold water (100 mL) and extracted with EtOAc (2 x 50 mL). The combined organic extracts were washed with brine solution (100 mL), dried over anhydrous $Na_2SO_4$, filtered, and concentrated under reduced pressure to obtain the crude. The crude was purified by flash column chromatography [eluent: 1% MeOH in DCM] to afford di-alkynylated meldrum acid (460 mg, 40% yield) as an off-white solid.

**Isolation of primary chondrocytes and culture**: Primary mouse chondrocytes were isolated from the articular cartilage of the knee joints obtained from the ~7-days-old C57BL-6 mice according to the previously mentioned protocol[1]. Briefly, the articular cartilage from the tibia and femur part was extracted from the knee joints of ~7-day-old C57BL-6 mice and dipped in serum-free DMEM (Life Technology, 10567014) solution at room temperature. After thorough washes with 1X PBS, cartilage tissues were incubated in the 3 mg/mL Collagenase D (Roche) in serum-free DMEM solution for 45-minute twice and then transferred to overnight incubation in 0.5 mg/mL Collagenase D in DMEM solution supplemented with 3% Liberase TL (Sigma). The next day, tissues were homogenized using pipettes to release and suspend the cells multiple times. Finally, the homogenate was filtered through a 40 μm strainer to remove large debris and resuspended in fresh DMEM supplemented 10% FBS and 1% Pen-strep solution. The suspension of cells was then plated in 60 mm cell culture dishes and allowed to expand as passage zero cells at 37 °C and 5% $CO_2$.

Other cell lines such as primary mouse adipocytes, and HEK-293T were also cultured in DMEM medium supplemented with 10% FBS and 1% Pen-strep solution.

**Metabolic labeling and cell lysis**: The cells were treated with MA-diyne as described in the main manuscript. The same volume of DMSO was taken as a negative control. After the stipulated incubation time for metabolic labeling, cells were washed with ice-cold 1X PBS thrice and scraped using a plastic scraper in 1X PBS. The cells were collected in a 1.5 mL centrifuge tube and centrifuged at 3000 rpm. The cell pellet thus obtained was lysed using ice-cold lysis buffer (1% NP 40, 150 mM NaCl, 50 mM HEPES, 2 mM $MgCl_2$, 20% Glycerol pH 7.4, EDTA free protease inhibitor cocktail) for 15 mins on ice followed by centrifugation at 16000g for 15 mins at 4°C. The supernatant was collected, and protein concentration was estimated using a BCA protein estimation kit (Pierce). The cell lysates were then diluted using dilution buffer (150 mM NaCl, 50 mM HEPES, 2 mM $MgCl_2$, 20% Glycerol pH 7.4, EDTA-free protease inhibitor cocktail) to give the final concentration of 1.5mg/mL for the click reaction.

**Copper (I) catalyzed alkyne-azide cycloaddition reaction (CuAAC)**: The procedure for CuAAC click chemistry reaction with protein lysates was performed as described in Yang et al 2010 with slight modification[2]. Briefly, a click reaction master mixture for the required volume of protein lysates was prepared by mixing the reagents in the following order: 100 µM IR680 azide dye (click chemistry tools) for in-gel fluorescence / 100 µM Biotin azide (Click chemistry tools) for streptavidin enrichment and western blotting, 100 µM Tris((1-benzyl-4-triazolyl)methyl)amine (TBTA) (Click chemistry tools), 1 mM Tris(carboxyethyl)phosphine (TCEP) (Sigma) and 1mM $CuSO_4$. The protein lysates were incubated with the click reaction master mix for 1.5 h at room temperature.

**In-gel fluorescence imaging of proteins**: After incubating the protein with click reaction mixture for 1.5 h, 4 volumes of ice-cold acetone were used to stop/quench the reaction. The proteins were then incubated at -20°C overnight to precipitate the proteins. The protein acetone solution was centrifuged at 6000 g for 5 min at 4°C. The obtained pellet was washed with ice-cold methanol once and then air-dried for 10 mins. The protein pellets were solubilized in 1X loading buffer with 100 mM DTT, heated at 95°C for 10 mins, and then resolved in 10 % SDS PAGE gel. The labeled proteins were visualized by scanning the gel under the Odyssey CLx Imaging System (excitation maxima-672 nm, emission maxima- 694 nm).

**Pulse-chase experiment**: Wild type primary chondrocytes and Sirt5KO cells were isolated from the wild type or the Sirt5 conditional knockout (Sirt5-CKO; Aggrecan-CreERT2; Sirt5 flox/flox) mice respectively. The primary chondrocytes cells (both wild type and SirtKO) were treated with 4-hydroxy tamoxifen for 48 h for inducing the cre-lox system. The cells were labeled with 200 µM MA-diyne for 1 h. The medium was removed, and the cells were washed thrice with 1X PBS to ensure removal of residual MA-diyne. Fresh medium supplemented with meldrum acid (200 µM) was added to the cells. The cells were extracted at different time points (0 to 5 h). The cells were then subjected to protein lysis as described earlier followed by click reaction with IR680 azide.

**Labeling the cells for confocal imaging:** The cells were cultured on sterile coverslips in 6 well culture dishes. The cells were treated with different concentrations of MA-diyne for 6 h and washed with 1X PBS. The cells were then fixed using ice-cold 4% paraformaldehyde solution for 20 mins at 4°C. The cells on coverslips were then washed thoroughly with ice-cold 1X PBS several times and permeabilized using 0.25% triton X-100 prepared in 1X PBS for 25 mins at room temperature. The cells were later washed with 1X PBS and then 5% BSA blocking buffer was added to the wells and left for shaking at room temperature for another 30 mins. The cells were then incubated with click reaction mix containing 100 µM Carboxyrhodamine 110 Azide, 100 µM TBTA, 1 mM TCEP and 1mM $CuSO_4$ in PBS for 1 h. Finally, cells were washed with 1X PBS to remove the remaining staining and coverslips were mounted on clean glass sclides with DAPI containing mounting reagent. The image was collected in Nikon A1R confocal microscope at 60X magnification.

**Measurement of Malonyl CoA in cell supernatant using malonyl CoA ELISA kit**: The cells were treated with MA-diyne for 2-4 h with different concentrations. After the incubation, cells were trypsinized and resuspended in 1X PBS at a concentration of 1 X $10^6$ cells/mL. The cells were lyzed by repeated freeze-thaw cycles and centrifuged at 2000-3000 rpm for 20 mins at 4°C. The supernatant was collected to estimate the malonyl CoA concentration using the mouse malonyl CoA ELISA kit (MyBioSource). The instructions for the experiment were followed as mentioned in the manufacturer's protocol.

**Pull down of labeled proteins after streptavidin enrichment:** The cells were labeled with MA-diyne as instructed above and then the protein lysates were conjugated with biotin azide via click reaction. The reaction was quenched using 4 volumes of ice-cold acetone and the solution was incubated at -20°C overnight to precipitate the proteins. The proteins were pelleted at 3000g for 5 mins and washed with methanol thrice. The protein pellet was air dried for 10 mins and solubilized in 100 µL solubilizing buffer (4% SDS, 20 mM EDTA, and 20 % glycerol in 1X PBS) by vortexing and gentle heating. The solution was diluted with 1X PBS to decrease the SDS concentration to 0.5 %. 100 µL of pre-washed (three times 0.2% SDS prepared in 1X PBS) avidin agar beads were added to the above protein solution and kept for incubation for 1.5 h at room temperature on a gyrator rocker. Centrifuge the solution at 3000 rpm for 2 mins and discard the supernatant. Wash the beads thoroughly as follows: 5 times with 10 mL 0.2% SDS in 1X PBS, 5 times with 10 mL 1X PBS, and finally 5 times with distilled water. Boil the beads in equal volume of 100 uL of elution buffer (loading buffer+ 100 mM DTT) at 95 °C for 5 mins and load the elute on the SDS PAGE gel. The proteins were then transferred to the nitrocellulose membrane and probed against anti streptavidin as well as the primary antibody of choice.

**Sample preparation for LC MS/MS :** The avidin beads bound to the biotinylated proteins were washed as instructed above and stored in -80°C before sending for the LC MS/MS at Cell Signaling Technology, Inc. Proteomics facility. The beads were washed again with HPLC water to prevent detergent or protease inhibitor contamination in the mass spectrometer. The beads were incubated with 9M urea for 10 mins at room temperature. The cysteines were then subjected to reductive alkylation with dithiothreitol (DTT) and iodoacetamide (IAA). Finally, the whole solution was diluted with 3 volumes of 20mM HEPES so that urea concentration reduces to less than 2M. The beads were then subjected to on-bead trypsin digestion overnight followed by addition of 1% TFA (final concentration). The peptides were then desalted using stagetip and eluted with 50% acetonitrile. Peptide concentration was measured using BCA assay and samples were dried. The enriched peptides was then subjected to bottom-up quantitative proteomics on Orbitrap Astral Instrument. Peptides were loaded directly onto an Aurora Ultimate TS column (25 cm x 75 µm ID, 1.7 µm C18) packed with C18 reversed-phase resin. The column was developed with a 45- minute linear gradient of acetonitrile at 400 nL/min. “A” buffer = 3% acetonitrile, 0.1% formic acid in water. “B” buffer = 80% acetonitrile, 0.1% formic acid in water. Full MS parameter settings are available upon request.

**Identification of target proteins**: MS/MS spectra were evaluated using Spectronaut software from Biognosys. Hybrid DIA searches were performed using factory default settings. Searches were performed against the most recent update of the Uniprot Mus musculus database. Tryptic cleavage was required with up to 4 missed cleavage sites per peptide allowed. Carbamidomethyl modification on cysteine was kept as static modification while oxidized methionine and N-terminal acetylation was set as variable modifications with maximum 5 modifications/peptide. All the results were filtered to a 1% false discovery rate at the precursor peptide level.

**Bioinformatics:** Gene Ontology analysis (https:// predictprotein.org/) was used for identify the sub-cellular localization and biological processes of the proteins enriched by MA-diyne. A Kyoto Encyclopedia of Genes and Genomes (KEGG) pathway analysis was also conducted to evaluate the pathways involved (https://www.genome.jp/kegg/pathway.).

Ethical approval for the animal experiments reported here was obtained from the Institutional Animal Care and Use Committee (IACUC) at Ohio University, IACUC protocol number is 20-H-014.

**<u>Scheme SF1</u>. <u>$^{1}$H NMR Spectral data MA-diyne probe</u>**

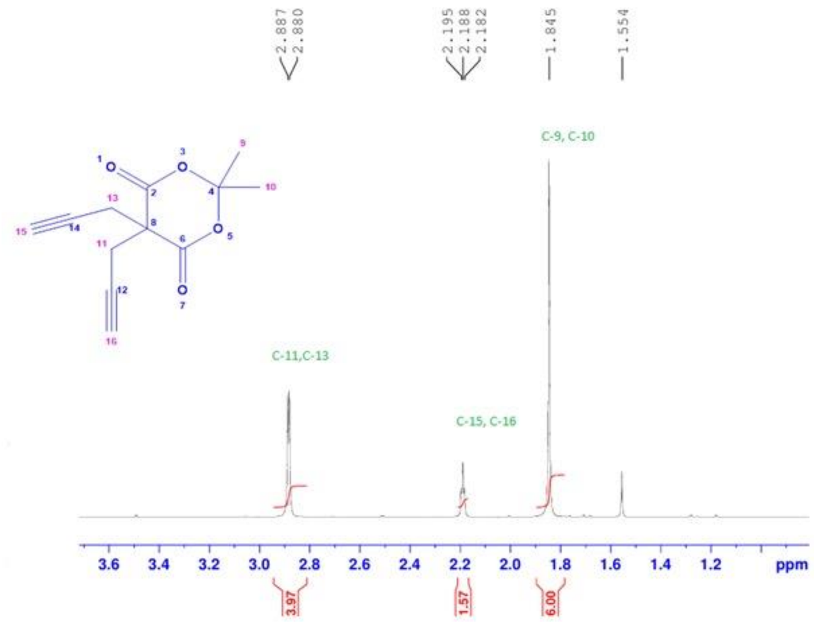


**Scheme SF2. <u>$^{1}$H NMR Full spectrum of MA-diyne probe:</u>** * TMS-internal standard; ▲-residual water in $CDCl_3$; #-$CDCl_3$ solvent)

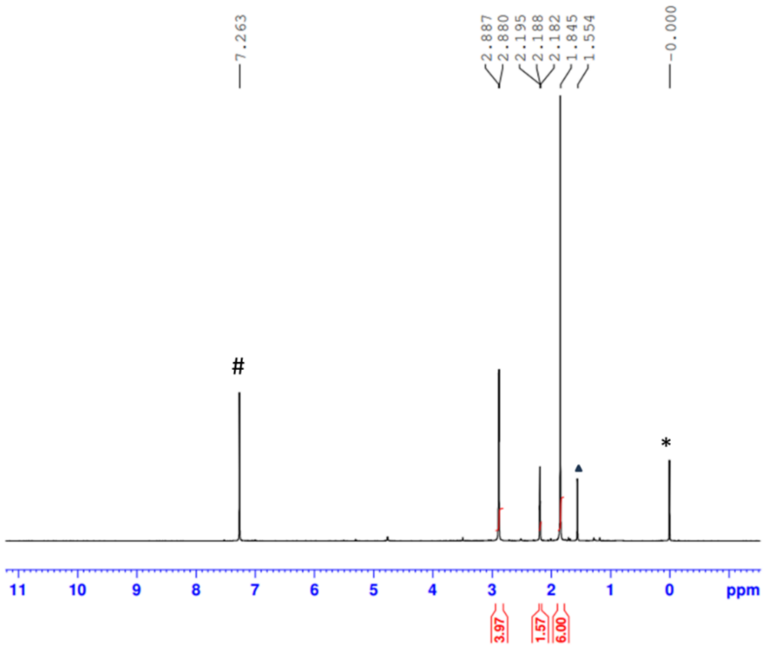

**Supplementary Figure 1**: Western blot analysis of the proteins obtained from primary chondrocytes treated with or without ND-630 [pan ACC (acetyl carboxylase CoA inhibitor)] using a commercial anti-KMal antibody.

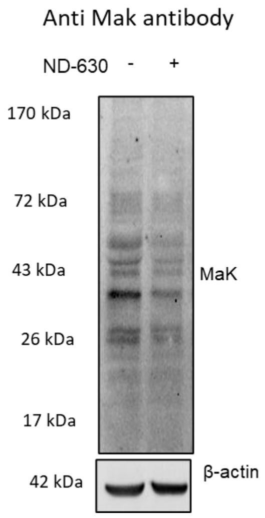

**Supplementary Figure 2**: Confocal microscopic fluorescence image of the cells treated with different concentrations of MA-diyne after click reaction with carboxyrhodamine 110 azide. Cells treated with vehicle/100 µM MA-diyne with and without click reaction were taken as control samples.

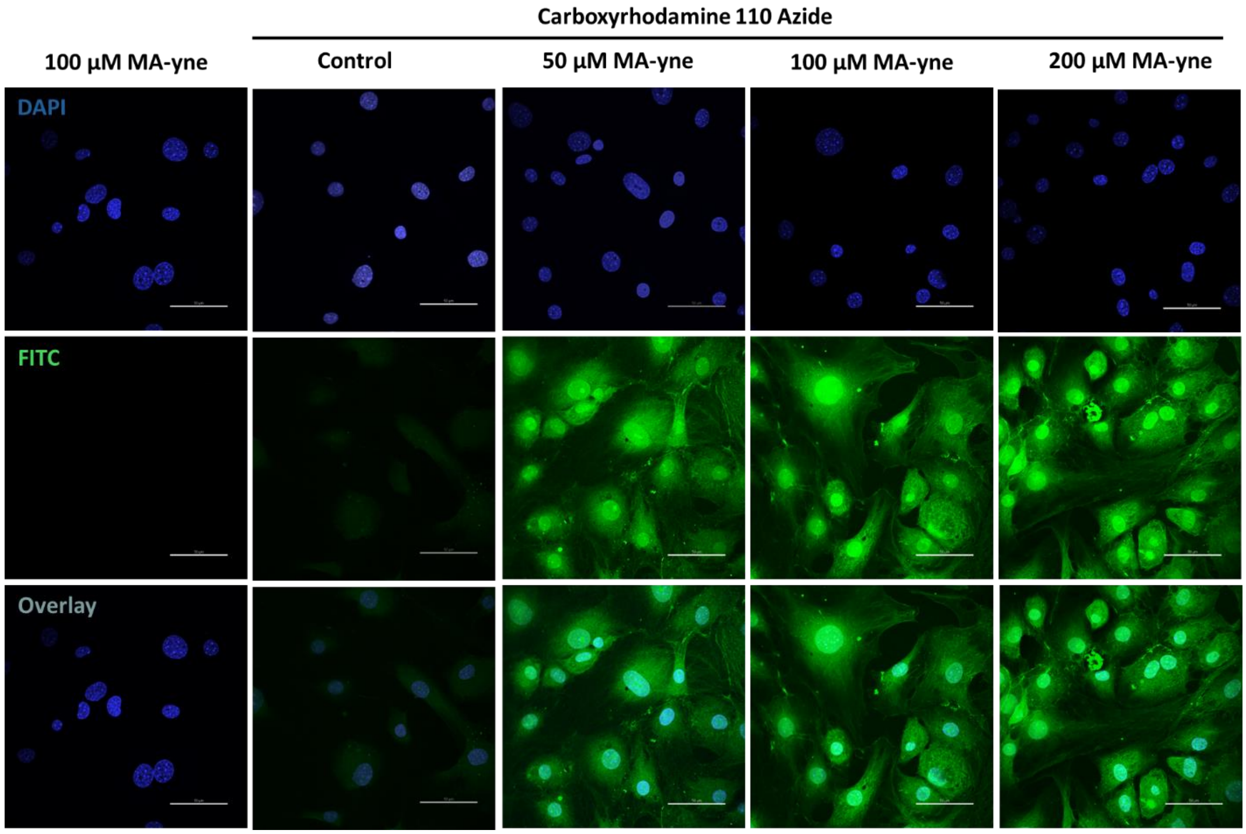

**Supplementary Figure 3**: Wild-type primary chondrocytes incubated with 10 nM ND-630 for 48 h and ACC1 KO primary chondrocytes were labeled with MA-diyne (100 µM) and the protein lysates were clicked with IR680 azide followed by in-gel fluorescence analysis.

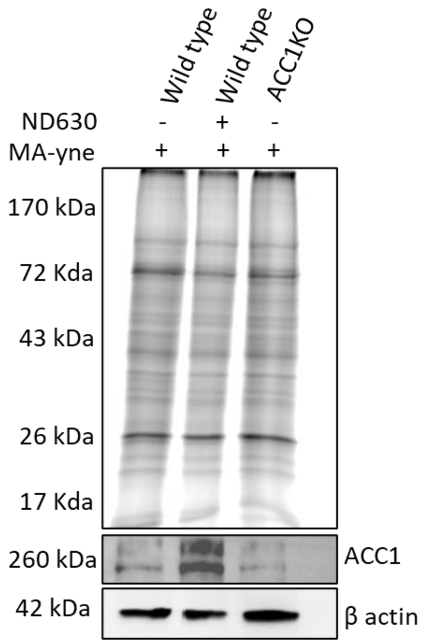


**References.**

1 Gosset, M., Berenbaum, F., Thirion, S. & Jacques, C. Primary culture and phenotyping of murine chondrocytes. *Nature Protocols* **3**, 1253-1260 (2008). https://doi.org/10.1038/nprot.2008.95

2 Yang, Y. Y., Grammel, M., Raghavan, A. S., Charron, G. & Hang, H. C. Comparative analysis of cleavable azobenzene-based affinity tags for bioorthogonal chemical proteomics. *Chem Biol* **17**, 1212-1222 (2010). https://doi.org/10.1016/j.chembiol.2010.09.012
